# A *Drosophila* model of chemotherapy-related cognitive impairment

**DOI:** 10.1101/2023.06.01.543297

**Authors:** Matthew Torre, Hassan Bukhari, Vanitha Nithianandam, Camila A Zanella, Douglas A Mata, Mel B Feany

**Author notes:** (Corresponding author) Mel B Feany, MD/PhD, Department of Pathology, Brigham and Women’s Hospital, 75 Francis Street Boston, MA 02115.

## Abstract

Chemotherapy-related cognitive impairment (CRCI) is a common adverse effect of treatment and is characterized by deficits involving multiple cognitive domains including memory. Despite the significant morbidity of CRCI and the expected increase in cancer survivors over the coming decades, the pathophysiology of CRCI remains incompletely understood, highlighting the need for new model systems to study CRCI. Given the powerful array of genetic approaches and facile high throughput screening ability in Drosophila, our goal was to validate a *Drosophila* model of CRCI. We administered the chemotherapeutic agents cisplatin, cyclophosphamide, and doxorubicin to adult Drosophila. Neurocognitive deficits were observed with all tested chemotherapies, especially cisplatin. We then performed histologic and immunohistochemical analysis of cisplatin-treated *Drosophila* tissue, demonstrating neuropathologic evidence of increased neurodegeneration, DNA damage, and oxidative stress. Thus, our *Drosophila* model of CRCI recapitulates clinical, radiologic, and histologic alterations reported in chemotherapy patients. Our new *Drosophila* model can be used for mechanistic dissection of pathways contributing to CRCI and pharmacologic screens to identify novel therapies to ameliorate CRCI.

**Summary Statement:** We present a *Drosophila* model of chemotherapy-related cognitive impairment, which recapitulates neurocognitive and neuropathologic changes observed in cancer patients treated with chemotherapy.

## Introduction

The number of cancer survivors in the United States currently exceeds 18 million (Miller, et al., 2022) and is expected to increase to 26 million by 2040 (Bluethmann, et al., 2016). Conventional chemotherapy remains a mainstay of cancer treatment, particularly for locally advanced or metastatic tumors. Correspondingly, the global demand for chemotherapy is estimated to rise by over 50% between 2018 and 2040 (Wilson, et al., 2019).

Up to 80% of chemotherapy patients develop cognitive deficits (Janelsins, et al., 2014), an observation known as chemotherapy-related cognitive impairment (CRCI) or, more colloquially, “chemobrain.” CRCI affects multiple cognitive domains including memory, attention, and executive function (Pendergrass, et al., 2018). The cognitive deficits observed following chemotherapy treatment are functionally significant and can contribute to reduced ability to function at work or return to work (Mehnert, 2011). CRCI may persist for decades after therapy (Koppelmans, et al., 2012), and some epidemiologic studies have found an association between chemotherapy history and subsequent risk of dementia (Chiu, et al., 2022; Heck, et al., 2008). Radiologic brain alterations following chemotherapy treatment have been reported, such as decreases in grey matter volume and density, functional/structural connectivity changes, and reduced white matter integrity (Bruno, et al., 2012; de Ruiter, et al., 2012; Li, et al., 2018; McDonald, et al., 2012; Niu, et al., 2021). Risk factors for developing CRCI include advanced age, low cognitive reserve, and certain genetic polymorphisms (e.g. APOE4 allele) (Ahles, et al., 2010; Mandelblatt, et al., 2018).

Despite the projected increase in cancer survivors and the impact of CRCI on patients’ quality of life and daily functioning (Boykoff, et al., 2009), the pathophysiology of CRCI is incompletely understood (Karschnia, et al., 2019; Ren, et al., 2019; Torre and Feany, 2020). In vivo CRCI models have been developed in rodents and have implicated mechanisms involving neuroinflammation, proinflammatory cytokines, oxidative stress, DNA damage, direct cytotoxicity on brain cell populations, dysmyelination, and neurogenesis, among others (Geraghty, et al., 2019; Gibson, et al., 2019; John, et al., 2021). Our goal was to validate a *Drosophila* model of CRCI that can be used as a complementary system to investigate the pathophysiology of CRCI.

*Drosophila* melanogaster, the common fruit fly, is a powerful system to study human disease. Approximately 75% of human disease-causing genes have *Drosophila* orthologs (Reiter, et al., 2001), with most having a conserved cellular function (Ugur, et al., 2016). Compared to mammalian model organisms, *Drosophila* have rapid generation times, shorter life cycles, and are inexpensive to grow and maintain. Importantly, use of *Drosophila* is highly amenable to forward and reverse genetic approaches, including gene editing, fast generation of transgenic lines with tissue-or cell type-specific expression of genes of interest, and high throughput pharmacologic and genetic modifier screens. The strengths of the *Drosophila* model system have resulted in important mechanistic insights into neurodegenerative and neurodevelopmental disorders and forms of iatrogenic neuropathology including chemotherapy-induced peripheral neuropathy (Bussmann and Storkebaum, 2017; Feany and Bender, 2000; Gatto and Broadie, 2011; Lu and Vogel, 2009), highlighting the importance and practicality of a *Drosophila* CRCI model.

To establish a *Drosophila* model of CRCI, we wanted to determine if chemotherapy-treated flies recapitulate the clinical and neuropathologic features of chemotherapy patients. Thus, we investigated whether chemotherapy-treated flies show neurocognitive deficits, evidence of neurodegeneration (corresponding to patients’ neuroradiology changes), and increased DNA damage and oxidative stress, which we have previously described in the brains of chemotherapy patients (Torre, et al., 2021).

## Results

### Chemotherapy-treated *Drosophila* show neurocognitive deficits

To investigate the effects of chemotherapy administration to flies, adults were aged to 5 days and then transferred to vials containing instant *Drosophila* medium rehydrated with chemotherapy-containing aqueous solution or vehicle (H_2_O) control. Flies were treated with a 3 day regimen of chemotherapy or vehicle, and then transferred back to vials containing standard cornmeal-agar medium and aged to 10, 20, or 30 days for neurocognitive testing (climbing assay, taste memory assay) or tissue harvesting. The chemotherapeutic regimens used were cisplatin (10, 100, or 500 μg/ml), cyclophosphamide (10, 100, or 1000 μg/ml), and doxorubicin (10, 100, or 1000 μg/ml).

The climbing assay was performed to assess general neurologic function and is based on the negative geotaxis of *Drosophila* (i.e. their intrinsic proclivity to climb upwards). Flies treated with chemotherapy and aged to 10 days showed reduced climbing ability (Figure 1). Comparisons with the vehicle control group reached statistical significance for the intermediate and high dose cisplatin cohorts (100 and 500 μg/ml), low and intermediate dose cyclophosphamide cohorts (10 and 100 μg/ml), and low, intermediate, and high dose doxorubicin cohorts (10, 100, and 1000 μg/ml). A trend towards reduced climbing ability for the chemotherapy cohorts was also observed at day 20. However, due to the low percent of aged control flies with intact climbing ability at day 20, the study was not sufficiently sensitive to detect statistically significant differences between groups.

**Figure 1.**
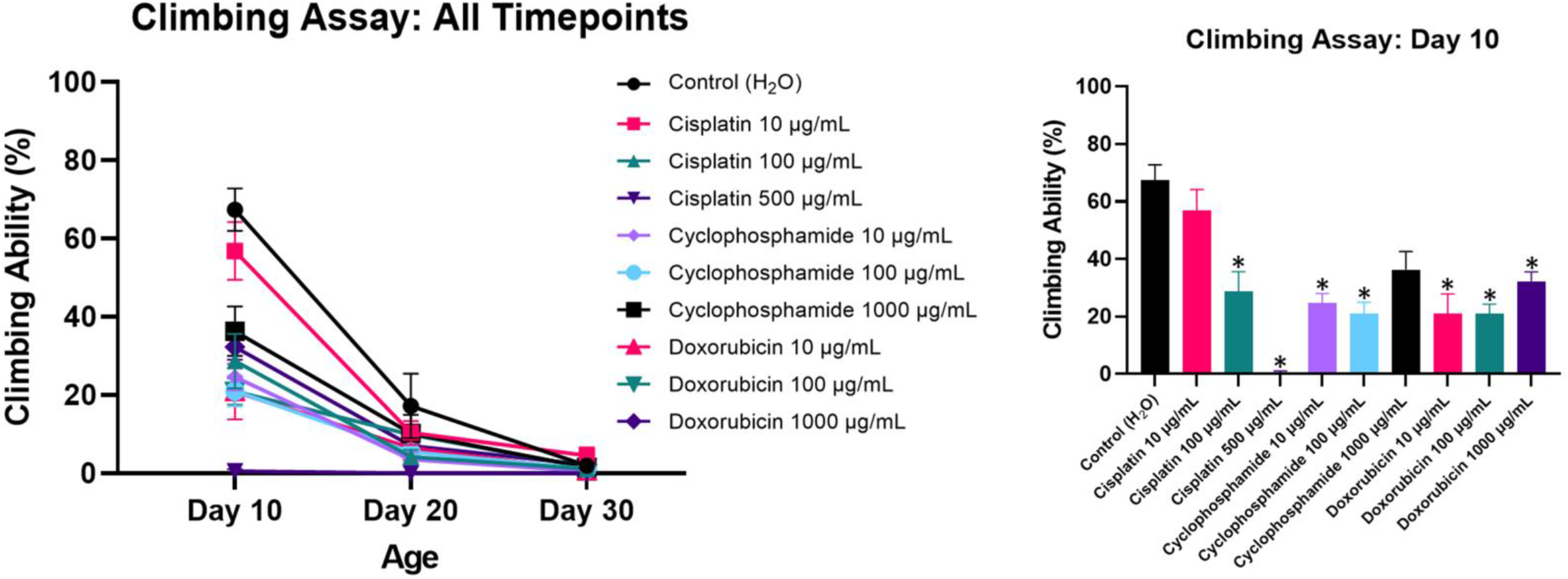
Chemotherapy-treated *Drosophila* show impaired climbing ability, a general neurologic readout. Statistically significant differences are observed at day 10. Data are represented as the mean ± SEM. Statistical analysis was performed using the Whitney-Mann U test and Bonferroni correction for multiple testing (corrected p value <0.00555 considered significant). Each cohort consists of n=6 replicates of 7-12 flies (≥53 flies total/cohort) at 10, 20, or 30 days post-eclosion. The genotype is *w*^*1118*^. *p value <0.00555; asterisks are included on the bar graph only.

The taste memory assay was performed to assess learning and memory. The taste memory assay is a negative reinforcement conditioning test and takes advantage of the *Drosophila* proboscis extension reflex (PER). When sucrose solution is presented to their tarsi, *Drosophila* will extend their proboscises to feed. During the taste memory assay, *Drosophila* are trained not to extend their proboscises when their tarsi are exposed to sucrose solution or else their labella are exposed to a bitter, aversive quinine solution. The assay evaluates the ability of flies to learn and remember not to extend their proboscises. Importantly, flies without an intact PER during the pretest phase of the assay (indicating impaired motor ability to extend the proboscis or attenuated taste sensation) are excluded from the study. Flies treated with chemotherapy had impaired learning and memory compared to vehicle control flies (Figure 2). Flies exposed to chemotherapy had a higher percent of trials with a positive proboscis extension reflex during multiple timepoints in the test phase, indicating a reduced ability to learn and retain memory of the aversive stimulus. Statistically significant differences in performance in the test phase were observed for the high dose cisplatin cohort at minute 0 and 45, the intermediate dose cisplatin cohort at minute 5, the low dose cisplatin cohort (10 μg/ml) at minute 5 and 10, and the high dose doxorubicin cohort at minute 5.

**Figure 2.**
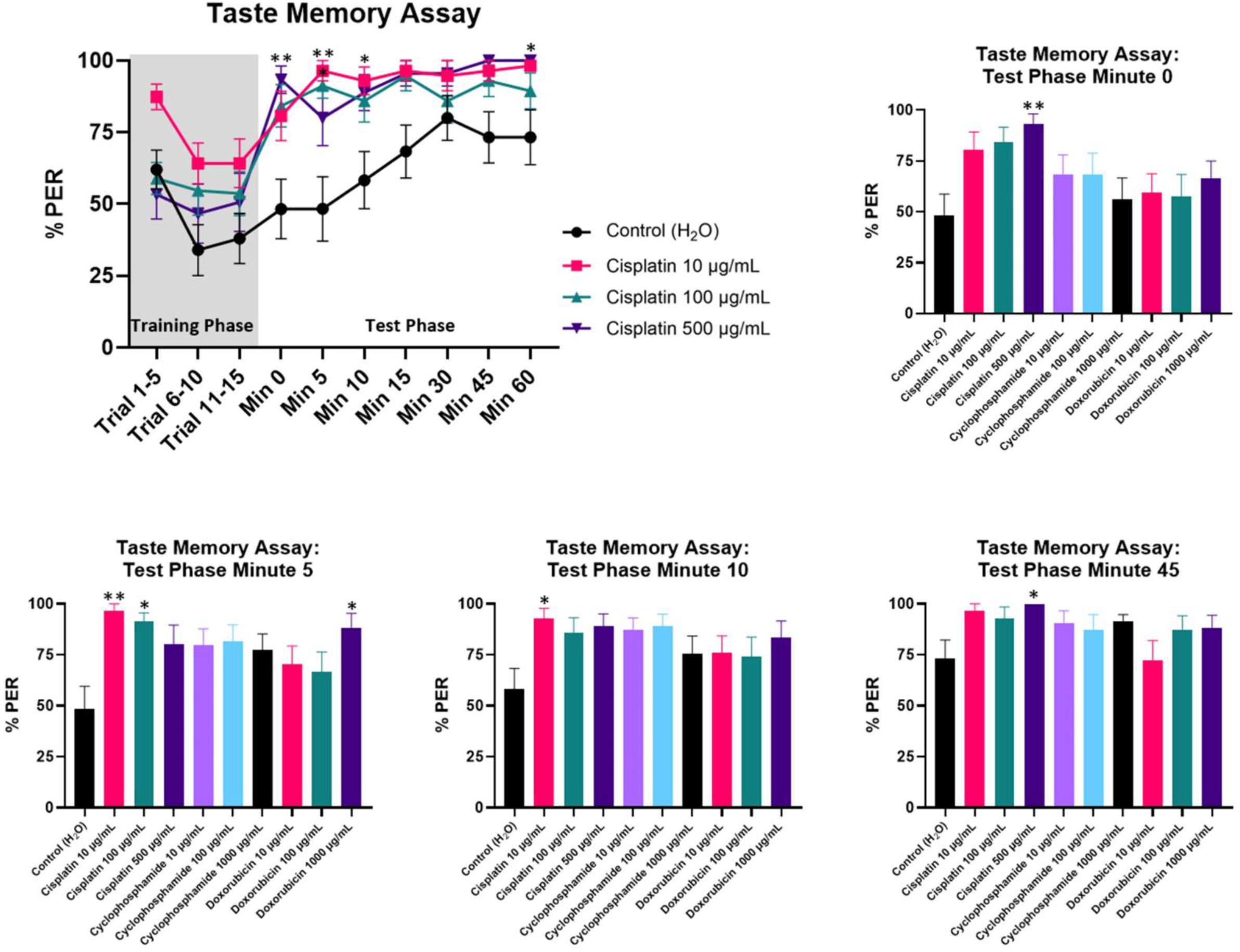
Chemotherapy-treated *Drosophila* have impaired learning and memory as assessed by the taste memory assay. Statistically significant differences are observed at minute 0, 5, 10, and 45 of the test phase for cisplatin- and doxorubicin-treated flies (see bar graphs). The line graph highlights differences in learning and memory between cisplatin and control cohorts. Data are represented as the mean ± SEM. Statistical analysis was performed using the repeated measures 2-way ANOVA with Dunnett’s multiple comparison test. Each cohort consists of n≥15 flies at 20 days post-eclosion. The genotype is *w*^*1118*^. *p<0.05, **p<0.01. PER, proboscis extension reflex.

Thus, chemotherapy-treated *Drosophila* demonstrate neurocognitive and behavioral deficits as assessed by the climbing assay and taste memory assay, suggesting that *Drosophila* will provide a useful model to study CRCI. The neurocognitive deficits are observed with different chemotherapeutic agents (over a range of concentrations) with diverse mechanisms of action.

### Cisplatin exposure exacerbates neurodegeneration in Drosophila

Because treatment with cisplatin resulted in the most severe neurocognitive deficits, we decided to focus our subsequent tissue analysis on flies given cisplatin. We assessed neurodegeneration in *Drosophila* by quantifying the number of cells with caspase activation and the number of brain vacuoles. We used da-GAL4 to drive ubiquitous expression of transgenic caspase reporter UAS-CD8-PARP-Venus. The UAS-CD8-PARP-Venus reporter flies contain a transgenic construct composed of the extracellular and transmembrane domains of mouse CD8 fused to a 40 amino acid sequence from human PARP that includes the caspase cleavage site (Williams, et al., 2006). Activated endogenous *Drosophila* caspases cleave human PARP at the caspase cleavage site. Caspase activation is then assessed using an antibody that specifically binds to cleaved human PARP (Bardai, et al., 2018; Frost, et al., 2014; Hegde, et al., 2014). We found significantly increased numbers of cells with caspase activation in the high dose cisplatin cohort at day 10, 20, and 30, the intermediate dose cisplatin cohort at day 30, and the low dose cisplatin cohort at day 10 compared to the vehicle control cohort (Figure 3). Double labeling immunofluorescence with elav, a neuron specific marker, demonstrated that the brain cells with activated caspase were predominantly neurons. Flies treated with high dose cisplatin also had significantly increased brain vacuoles compared to vehicle control at day 30 (Figure 4). The formation of brain vacuoles is frequently associated with neurodegeneration in *Drosophila* (Buchanan and Benzer, 1993; Katzenberger, et al., 2013; Ordonez, et al., 2018; Wittmann, et al., 2001). In summary, exposure to chemotherapeutic agent cisplatin was associated with elevated indicators of neurodegeneration, including caspase activation and brain vacuoles.

**Figure 3.**
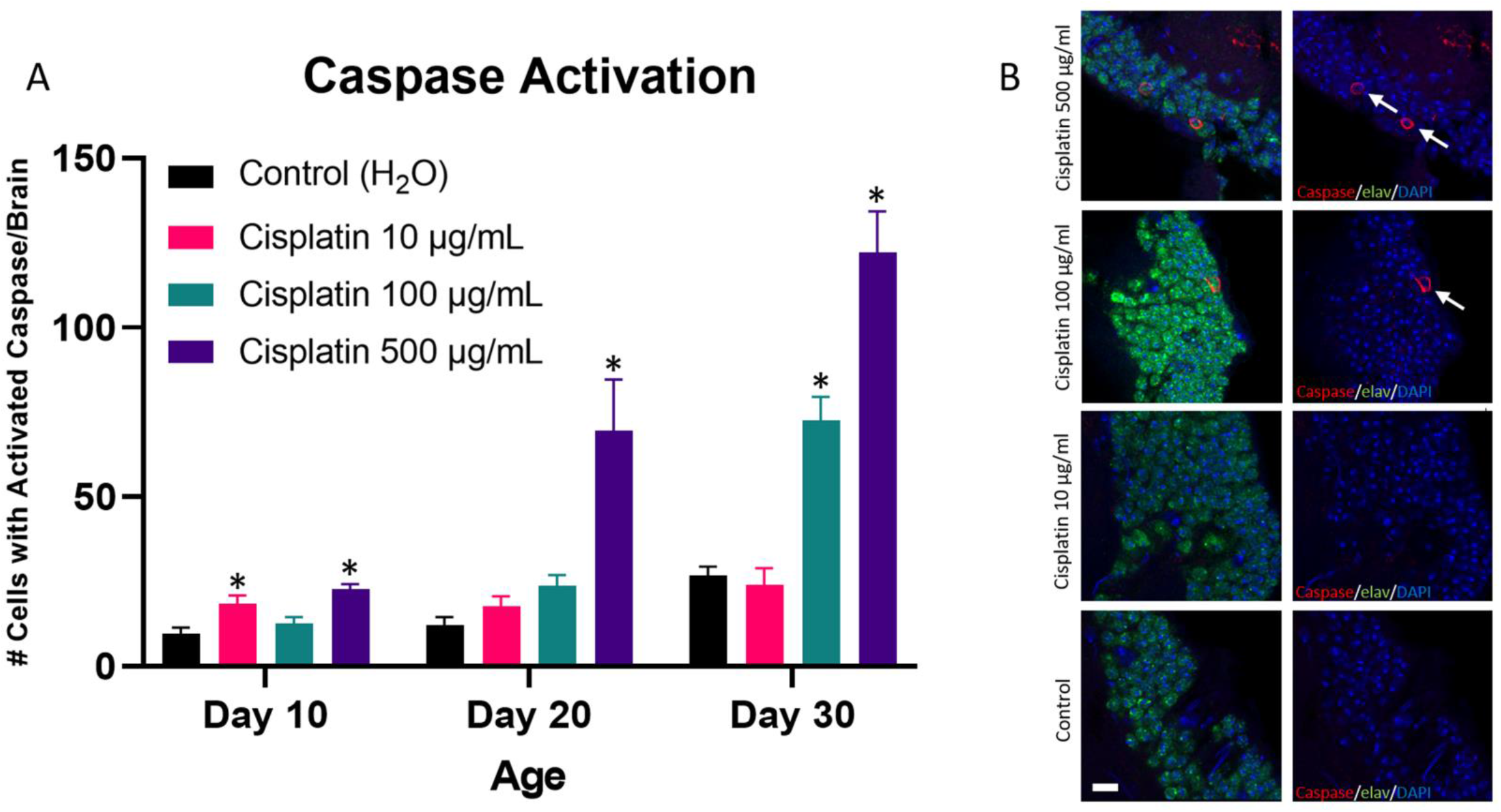
Neurodegeneration is increased in cisplatin-treated Drosophila, as assessed by caspase activation. (A) Flies given cisplatin have an increase in the number of brain cells with caspase activation, predominantly elav positive neurons. (B) Immunofluorescence images of brains from cisplatin and control flies at day 30 are shown (scale bar, 5 μm). Arrows highlight neurons showing caspase activation. Data are represented as the mean ± SEM. Statistical analysis was performed using the Whitney-Mann U test and Bonferroni correction for multiple testing (corrected p value <0.0166 considered significant). N=6 for all cohorts at 10, 20, and 30 days post-eclosion. The genotype is *UAS-CD8-PARP-Venus / da-GAL4*. *p<0.0166.

**Figure 4.**
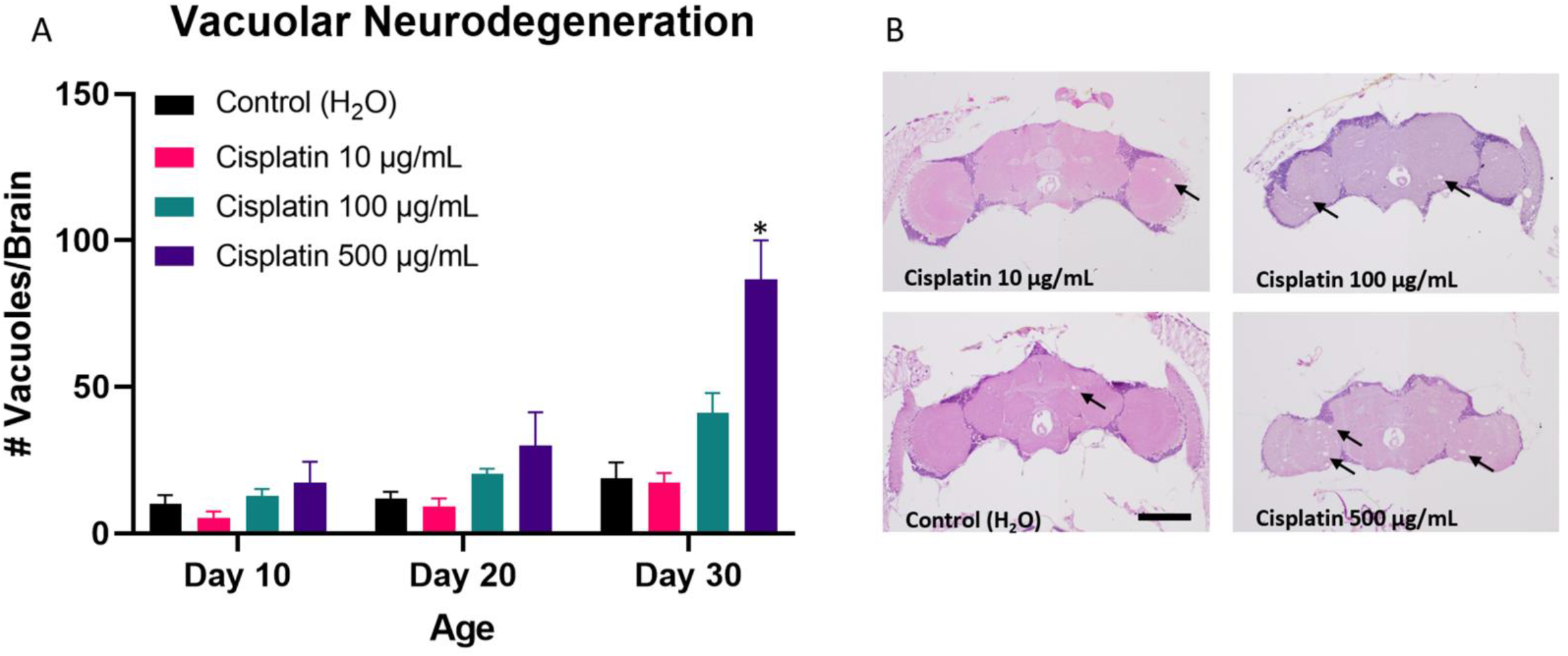
Neurodegeneration is increased in cisplatin-treated Drosophila, as assessed by brain vacuoles. (A) Flies given cisplatin have an increase in the number of brain vacuoles. (B) Hematoxylin and eosin images of brains from cisplatin and control flies at day 30 are shown (scale bar, 100 μm). Arrows highlight representative vacuoles. Data are represented as the mean ± SEM. Statistical analysis was performed using the Whitney-Mann U test and Bonferroni correction for multiple testing (corrected p value <0.0166 considered significant). N=6 for all cohorts at 10, 20, and 30 days post-eclosion. The genotype is *w*^*1118*^. *p<0.0166.

### DNA damage is elevated after cisplatin exposure

DNA damage was assessed by quantifying the percent of cells showing phosphorylation of serine 137 of histone variant H2Av (pH2Av), a marker of DNA double-strand breaks. We focused our quantification on an anatomically consistent region of the mushroom body, an area of the *Drosophila* brain that is important for learning and memory (Fiala, 2007; Zars, et al., 2000). A significantly increased percent of pH2Av positive cells was seen in the high dose cisplatin cohort at day 10, 20, and 30 and the intermediate dose cisplatin cohort at day 10 and 30 compared to the vehicle control cohort (Figure 5). Double labeling immunofluorescence with the neuronal marker elav demonstrated that the pH2Av positive brain cells were predominantly neurons. The results show that chemotherapy exposure results in increased DNA damage in the brain.

**Figure 5.**
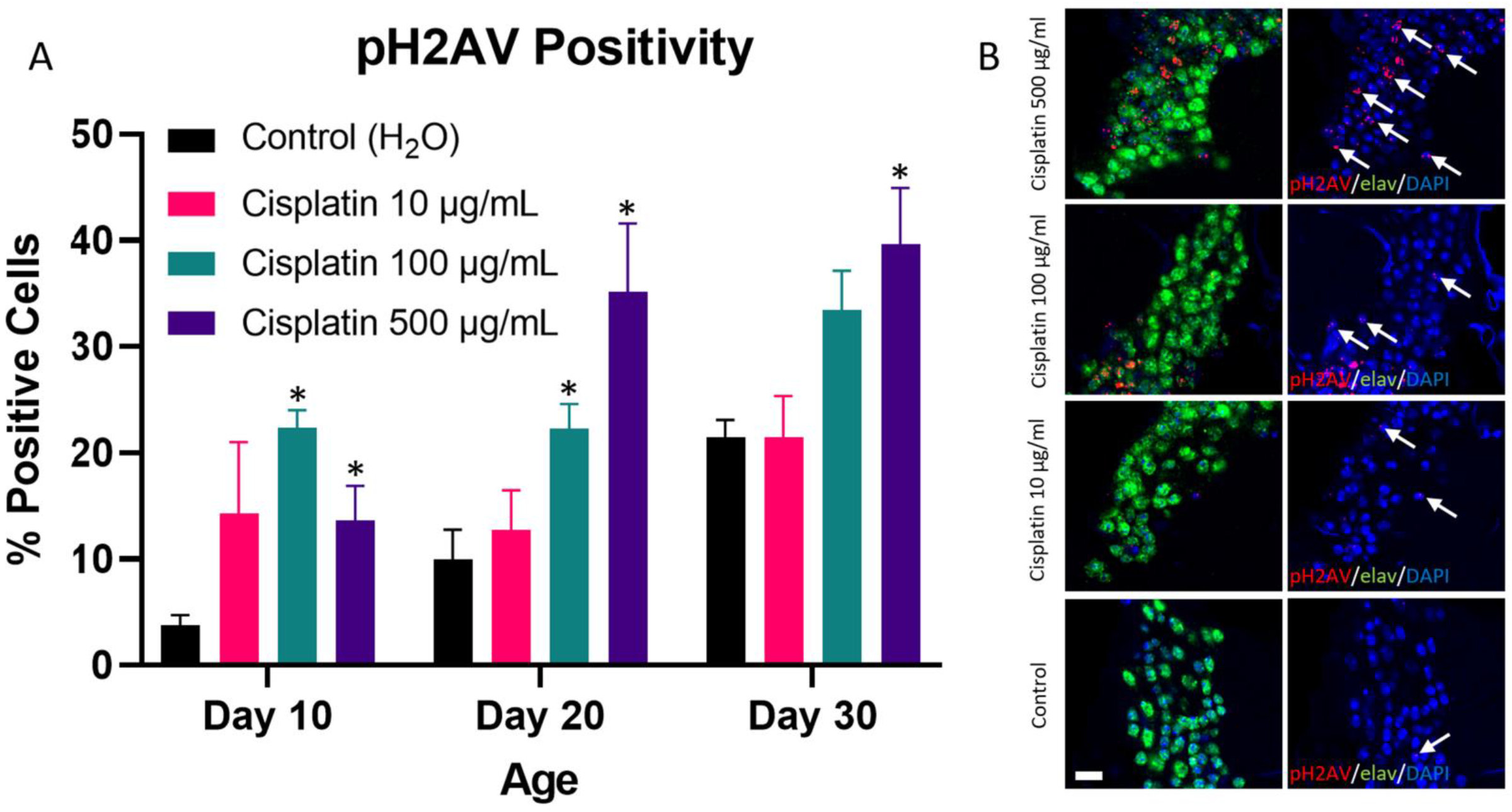
DNA damage is increased in cisplatin-treated Drosophila. (A) Flies given cisplatin have an increase in the percent of pH2Av positive cells, predominantly elav positive neurons, within an anatomically consistent region of the mushroom body. (B) Immunofluorescence images of brains from cisplatin and control flies at day 30 are shown (scale bar, 5 μm). Arrows highlight representative neurons with pH2Av punctate staining. Data are represented as the mean ± SEM. Statistical analysis was performed using the Whitney-Mann U test and Bonferroni correction for multiple testing (corrected p value <0.0166 considered significant). N=6 for all cohorts at 10, 20, and 30 days post-eclosion. The genotype is *w*^*1118*^. *p<0.0166.

### Oxidative stress is elevated after cisplatin exposure

Oxidative stress was assessed using the puckered-lacZ (puc-lacZ) and GstD1-GFP reporter systems. The puc gene encodes a phosphatase that is both a downstream target and a negative regulator of the JNK signaling pathway, which is activated in response to oxidative stress (Dias-Santagata, et al., 2007; Martin-Blanco, et al., 1998; McEwen and Peifer, 2005). Activation of the puc-lacZ reporter can be detected by β-galactosidase immunopositivity. We found that flies treated with cisplatin had increased numbers of brain cells with puc-lacZ activation compared to flies treated with vehicle control, reaching statistical significance in the high dose cisplatin cohort at day 30 and the intermediate dose cisplatin cohort at day 20 and 30 (Figure 6). Double labeling immunofluorescence with elav (data not shown) demonstrated puc-lacZ activation in neurons, as previously reported (Dias-Santagata, et al., 2007). GstD1 is an oxidative stress response gene and encodes glutathione S-transferase D1 (Wang, et al., 2003). Activation of the GstD1-GFP reporter (Sykiotis and Bohmann, 2008) can be detected by GFP immunopositivity. Preliminary immunofluorescence data suggested that differential activation of GstD1-GFP was more prominent in the retina than the brain following chemotherapy. Subsequent analysis confirmed that cisplatin-treated flies had increased cells in the retina with GstD1-GFP activation compared to vehicle control, reaching statistical significance in the high dose cisplatin cohort at day 20 and the intermediate dose cisplatin cohort at day 30 (Figure 7). In summary, chemotherapy exposure was associated with elevated markers of oxidative stress.

**Figure 6.**
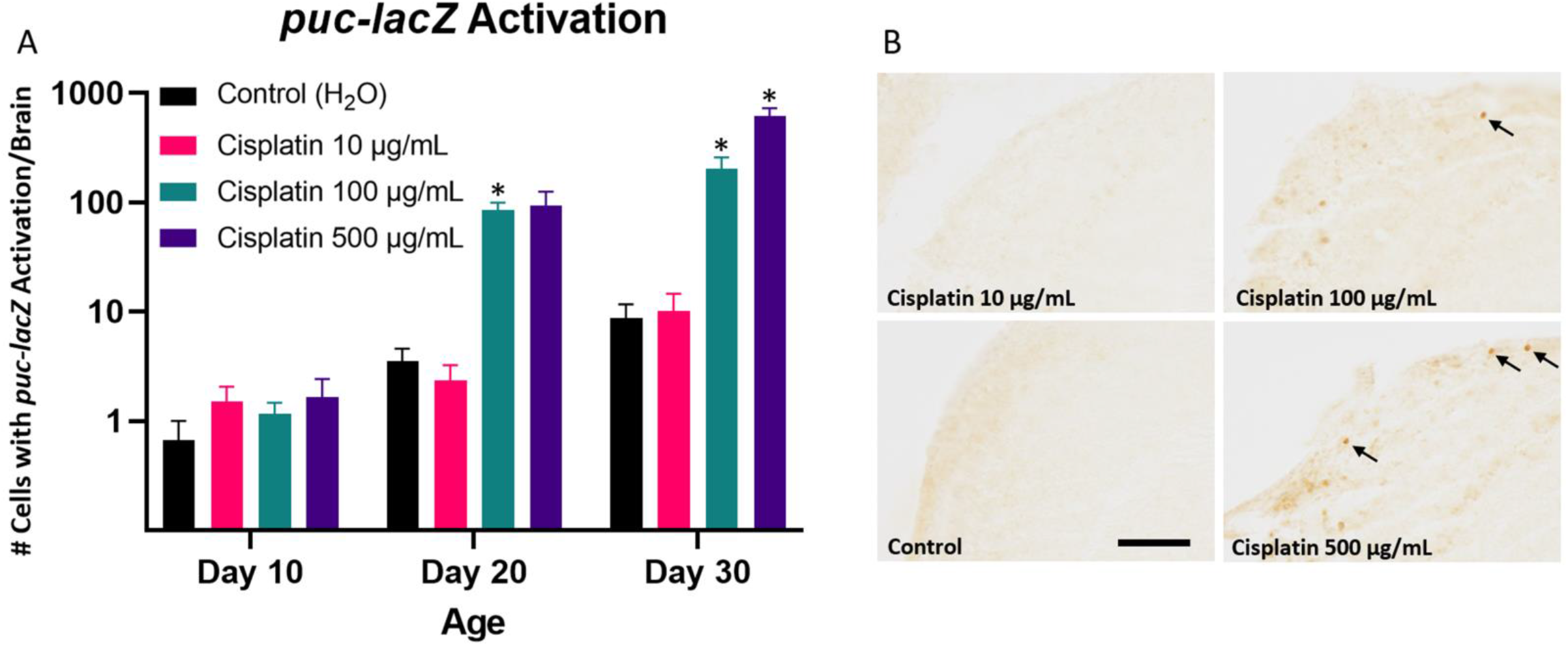
Oxidative stress is increased in cisplatin-treated Drosophila, as assessed by *puc-lacZ* activation. (A) Flies given cisplatin have increased brain cells with *puc-lacZ* reporter activation, indicated by β-galactosidase immunopositivity. (B) Immunohistochemical images of brains from cisplatin and control flies at day 30 are shown (scale bar, 20 μm). Arrows highlight representative cells with puc-lacZ activation. Data are represented as the mean ± SEM. Statistical analysis was performed using the Whitney-Mann U test and Bonferroni correction for multiple testing (corrected p value <0.0166 considered significant). N=6 for all cohorts at 10, 20, and 30 days post-eclosion. The genotype is *puc-lacZ/+*. *p<0.0166.

**Figure 7.**
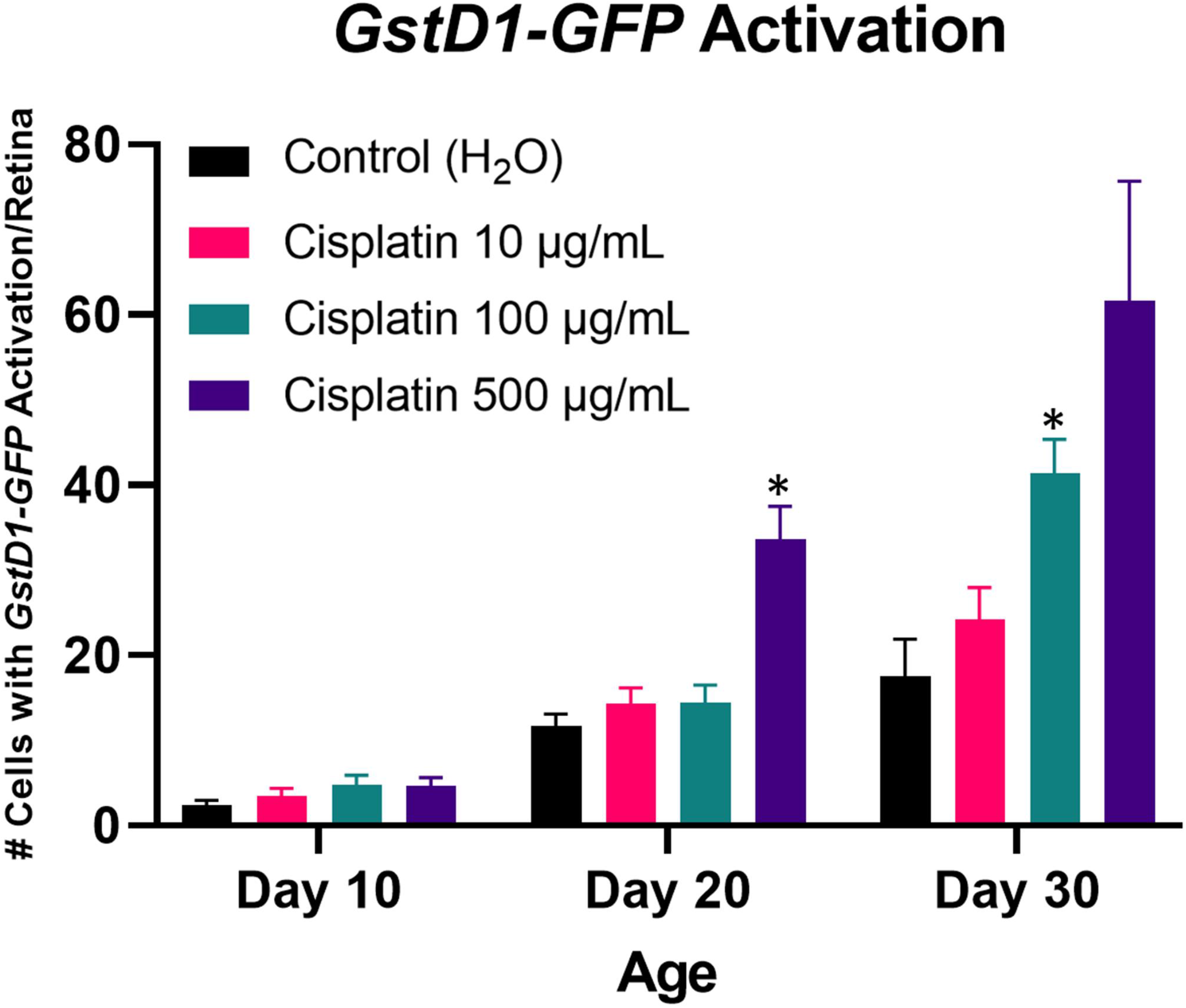
Oxidative stress is increased in cisplatin-treated Drosophila, as assessed by *GstD1-GFP* activation. Flies given cisplatin demonstrate an increased number of retinal cells with *GstD1-GFP* reporter activation. Data are represented as the mean ± SEM. Statistical analysis was performed using the Whitney-Mann U test and Bonferroni correction for multiple testing (corrected p value <0.0166 considered significant). N=6 for all cohorts at 10, 20, and 30 days post-eclosion. The genotype is *GstD1-GFP/+*. *p<0.0166.

## Discussion

CRCI is a clinically important and functionally significant adverse effect of chemotherapy observed in up to 80% of patients (Janelsins, et al., 2014). However, development of effective CRCI disease-modifying therapies is hindered by a limited understanding of the pathophysiology of CRCI. Thus, new approaches to investigate CRCI are needed. We present a working *Drosophila* model of CRCI that recapitulates clinical and neuropathologic features described in human patients.

Our *Drosophila* CRCI model shows neurocognitive deficits analogous to the clinical phenotype of chemotherapy patients. The climbing assay is a general neurologic readout that can be used as a high throughput screening test in young adult flies. The taste memory assay evaluates learning and memory and is thus particularly well-suited to model CRCI. Importantly, unlike other methods of assessing cognition in *Drosophila* such as the olfactory conditioning assay or aversive phototaxis suppression assay, the taste memory assay is not dependent on locomotor ability.

We observed neurocognitive deficits in our *Drosophila* model following treatment with different chemotherapeutic agents (cisplatin, cyclophosphamide, and doxorubicin) that have diverse mechanisms of action. Cisplatin, cyclophosphamide, and doxorubicin are highly relevant to CRCI. These chemotherapeutic agents are commonly administered to breast cancer patients, the patient population most studied in CRCI (Van Dyk and Ganz, 2021), and are among a group of chemotherapies whose use is associated with CRCI (John, et al., 2021; Lange, et al., 2019). The concentrations of chemotherapy that we tested were informed by previously published doses administered to *Drosophila* (Costa and Nepomuceno, 2006; Groen, et al., 2022; Groen, et al., 2018; Guerra-Santos, et al., 2016; Lehmann, et al., 2003; Podratz, et al., 2013; Podratz, et al., 2011b; Stoffel, et al., 2020; Teixeira da Silva, et al., 2021; Zijlstra and Vogel, 1989). Our concentration of cisplatin ranged from 10 μg/ml to 500 μg/ml, and a prior study reported that a cisplatin regimen of 100 ug/ml resulted in a whole *Drosophila* body concentration comparable to that used in human patients and rodent models of cisplatin-induced neurotoxicity (Podratz, et al., 2011b). Cisplatin resulted in the most marked neurocognitive deficits in our model, so we decided to focus our neuropathologic evaluation on cisplatin-treated flies.

Given that chemotherapy patients show radiologic changes in the brain including reductions in grey matter volume and density (Li, et al., 2018; Niu, et al., 2021), we were interested in assessing neurodegeneration in chemotherapy-treated Drosophila. We demonstrated that the number of cells with caspase activation is increased after cisplatin exposure. Our finding is in keeping with previous studies showing increased apoptosis and cell death in vulnerable brain cell populations following chemotherapy in in vivo and in vitro models (Dietrich, et al., 2006; Scholz, et al., 2022; Yang, et al., 2011). We observed that the number of cells with activated caspase is elevated acutely after cisplatin treatment (i.e. at day 10) and remains persistently elevated at day 20 and 30. Thus, chemotherapy appears to induce long term changes that promote ongoing caspase activation with age. This is supported by our vacuolar neurodegeneration data, which showed an increase in brain vacuoles in the high dose cisplatin group at day 30 but not at earlier timepoints. The presence of brain vacuoles in the cortex and neuropil plausibly reflects cell death and loss of axons (Kretzschmar, et al., 1997; Min and Benzer, 1997; Wittmann, et al., 2001). Overall, our *Drosophila* CRCI model shows increased neurodegeneration by both the number of cells with caspase activation in the brain and by the number of brain vacuoles.

We have previously reported that chemotherapy patients have elevated DNA damage and oxidative stress in frontal lobe cortical neurons compared to control patients (Torre, et al., 2021). We thus wanted to assess whether these pathways were upregulated in our *Drosophila* CRCI model. Flies treated with cisplatin show increased DNA damage, indicated by pH2Av positivity, compared to vehicle control. Similar to our observations with caspase activation, DNA damage increases acutely after cisplatin treatment and continues to be elevated long after cessation of chemotherapy. Oxidative stress (indicated by *puc-lacZ* and *GstD1-GFP* reporter activation) is elevated in cisplatin-treated flies compared to vehicle control, but statistically significant differences are observed only at day 20 and 30. However, we cannot exclude the possibility that other markers of oxidative stress may be elevated more acutely.

Differences that we observed in our *Drosophila* model in the timing of DNA damage and oxidative stress may implicate certain biological pathways in chemotherapy-induced neurotoxicity in the brain. Cisplatin is known to damage both nuclear and mitochondrial DNA (Podratz, et al., 2011a), and our data suggest that DNA damage occurs early after chemotherapy exposure. Cisplatin-induced DNA damage promotes the production of mitochondrial derived reactive oxygen species (ROS) (Kleih, et al., 2019) that over time can result in measurable increases in oxidative stress. These ROS may cause additional DNA damage, creating a positive feedback loop that contributes to persistently elevated DNA damage, oxidative stress, and caspase activation long after the initial chemotherapy insult is removed. It is compelling to speculate whether cellular senescence or anastasis arising in the setting of chemotherapy may have a role in these persistent changes.

Our study has a few limitations. Firstly, while the data implicate promising molecular pathways in CRCI, more mechanistic analysis is needed to determine if the neurotoxic effects of chemotherapy are due to its direct action on brain cell populations or through an indirect mechanism or a combination of the two. Secondly, while we demonstrate that multiple chemotherapeutic agents cause neurocognitive deficits in Drosophila, our tissue analysis focused on cisplatin since it resulted in the most significant decrement to neurocognitive function. Although we anticipate that the use of *Drosophila* will accelerate the mechanistic understanding of CRCI, we should emphasize that the pathophysiology of CRCI is multifactorial and that *Drosophila* may better recapitulate some mechanisms than others. Blood brain barrier permeability to chemotherapeutic agents, alterations in neurogenesis, and microglial activation likely contribute to CRCI, but additional studies are needed to determine whether these mechanisms can be appropriately modeled in Drosophila.

In summary, we present a *Drosophila* model of CRCI. Our *Drosophila* model demonstrates neurocognitive deficits, increased neurodegeneration, and elevated DNA damage and oxidative stress, recapitulating the clinical and neuropathologic features of chemotherapy patients. Our *Drosophila* model may thus serve as a complementary system to existing rodent CRCI models and take advantage of powerful *Drosophila* genetic tools, high throughput screening, low operational cost, and short lifespan. Our model can be easily adapted to test other chemotherapeutic drugs at different concentrations, timepoints, and combinations. Future *Drosophila* CRCI studies can leverage forward and reverse genetic approaches to decipher molecular pathways contributing to CRCI, use high throughput screens to identify pharmacologic agents that may ameliorate CRCI, and investigate the mechanistic interactions between CRCI and tau pathology, synuclein pathology, cellular senescence, and anastasis, among other pathways relevant to neurodegeneration.

## Materials and Methods

### *Drosophila* stocks and genetics

All *Drosophila* crosses were performed at 25°C. The following stocks were obtained from the Bloomington *Drosophila* Stock Center (NIH P40OD018537) at Indiana University, Bloomington, IN: *w*^*1118*^, *da-GAL4*, and *puc*^*E69*^ (*puc-lacZ*). Darren Williams (Kings College, London, United Kingdom) provided the UAS-CD8-PARP-Venus stock. Dirk Bohmann (University of Rochester Medical Center, Rochester, NY) provided the GstD1-GFP stock. Male and female flies were used for all experiments.

### Chemotherapy regimens and maintenance

Flies of the appropriate genotype were aged to 5 days post-eclosion in vials containing standard cornmeal-agar medium and then transferred to vials of instant medium (Carolina Biological; Burlington, NC) rehydrated with chemotherapy-containing aqueous solution or with vehicle (H_2_O) control (Milli Q water, MilliporeSigma; Burlington, MA). Flies were placed on a 3 day regimen of rehydrated instant medium, which was replaced daily. Following this 3 day regimen, flies were transferred back to vials containing standard cornmeal-agar medium and then aged to 10, 20, or 30 days post-eclosion for testing and/or harvesting for histologic studies. Chemotherapeutic agents cisplatin (10, 100, and 500 μg/ml), doxorubicin (10, 100, and 1000 μg/ml), and cyclophosphamide (10, 100, and 1000 μg/ml) were dissolved in Milli Q water. Chemotherapeutic agents were purchased from MilliporeSigma. Drug concentrations for each chemotherapeutic agent are similar to those used in prior *Drosophila* studies (Costa and Nepomuceno, 2006; Groen, et al., 2022; Groen, et al., 2018; Guerra-Santos, et al., 2016; Lehmann, et al., 2003; Podratz, et al., 2013; Podratz, et al., 2011b; Stoffel, et al., 2020; Teixeira da Silva, et al., 2021; Zijlstra and Vogel, 1989). Aging was performed at 25°C.

### Climbing assay

Flies underwent the climbing assay at 10, 20, or 30 days post-eclosion. Individual flies were tested at one timepoint only. The climbing assay was performed based on standard protocols (Ordonez, et al., 2018). For each timepoint and treatment cohort, 7-12 flies were placed in individual empty vials for a total of 6 vials. At least 53 flies were tested in each group. Vials were gently tapped to bring the flies to the bottom of the vial, and the percent of flies that climbed above 5 cm within 10 seconds was recorded. The climbing assay was repeated 3 times for each vial of flies and averaged.

### Taste memory assay

The taste memory assay was performed on chemotherapy or vehicle control treated flies at 20 days post-eclosion and based on standard protocols with minor modifications (Cevik and Erden, 2012; Keene and Masek, 2012; Poudel and Lee, 2018). Flies were starved for 24 hours prior to testing by being transferred to empty vials with Milli Q water-soaked Kimwipes (Kimberly-Clark; Franklin, MA). Flies were then briefly anesthetized with CO_2_, fixed to glass slides with nail polish, and moved to a 25°C humidified incubator for a 3 hour recovery period. The slides were then vertically mounted and placed under a dissecting microscope to observe the fly behavior. Flies were satiated with purified water, and flies that are not satiated after 5 minutes were excluded from the study.

The taste memory assay consists of 3 phases: a pretest phase, a training phase, and a test phase. During the pretest phase, a 500mM sucrose solution was presented to the fly tarsi to confirm an intact proboscis extension reflex (PER). Flies with negative PER were excluded from further study. The training phase consists of 15 trials. During these trials, fly tarsi were exposed to 500 mM sucrose solution, but when the proboscis extended, a bitter, aversive solution (50 mM quinine) was presented to the labellum. Flies were allowed to drink the quinine solution for up to 2 seconds or until they retracted their proboscises. Between trials, the tarsi and labellum were washed with water, and the flies were allowed to drink fresh water to satiation. The training trials were binned into 3 groups of 5 trials (trials 1-5, 6-10, and 11-15). During the test phase, the 500 mM sucrose solution was presented to the tarsi at multiple time intervals (0, 5, 10, 15, 30, 45, and 60 minutes) but the labellum was not exposed to quinine solution if the proboscis extended. At each test timepoint, flies underwent 3 trials with a 10 second interval between trials. The percent of trials with a positive PER was recorded for each fly at each timepoint. After each test timepoint, the tarsi were washed with water, and the flies were allowed to drink fresh water to satiation. At the end of the test phase, flies were given sucrose solution to confirm an intact PER, and flies with negative PER were excluded from the analysis. Sucrose and quinine solutions were presented to the flies as droplets at the end of 1 mL syringes with 21 gauge needles. At least 15 flies were tested per cohort.

### Histology, immunohistochemistry, and immunofluorescence

Flies were harvested at 10, 20, or 30 days post-eclosion and fixed in formalin. Fly heads were embedded in paraffin and serially sectioned through the entire brain and retina (either 2 or 4 µm thick sections) and processed through a series of xylenes, ethanol, and water. Vacuolar neurodegeneration was evaluated in 2 µm brain sections stained with hematoxylin and eosin (H&E). For the immunostaining studies (4 µm sections), incubation with the following primary antibodies was performed following routine heat antigen retrieval (10 mM sodium citrate buffer, pH 6.0): anti-cleaved PARP (Abcam, E51, Cambridge, MA; 1:50,000 for immunohistochemistry (IHC), 1:500 for immunofluorescence (IF)), anti-β-galactosidase (Promega, Z3783, Madison, WI; 1:500 for IHC, 1:100 for IF), anti-GFP (NeuroMab, N86/8, Davis, CA; 1:500 for IHC and IF), anti-pH2Av (Rockland, 600-401-914, Pottstown, PA; 1:2500 for IHC, 1:1000 for IF), and anti-elav (Developmental Studies Hybridoma Bank, clones 9F8A9 and 7E8A10, Iowa City, IA; 1:5 (9F8A9) or 1:50 (7E8A10) for IF). For IHC, slides were then incubated with biotinylated secondary antibodies (Southern Biotech, Birmingham, AL; 1:200), and the markers were visualized with the avidin-biotin complex (ABC) detection system (Vector Laboratories, Burlingame, CA) and 3,3⍰-diaminobenzidine (DAB). pH2Av IHC slides were counterstained with hematoxylin. For IF, slides were incubated with Alexa Fluor-conjugated secondary antibodies (Invitrogen, Alexa 488, Alexa 555, Carlsbad, CA; 1:200) and mounted with 4’,6-diamidino-2-phenylindole (DAPI)-containing Fluoromount medium (Southern Biotech).

For quantification of vacuolar neurodegeneration, caspase activation, and puc-lacZ activation, the number of vacuoles or the number of immunopositive cells throughout the entire brain was counted per fly, and the average number of vacuoles or immunopositive cells per brain was averaged for each cohort (n=6 per cohort). We quantified *GstD1*-*GFP* activation by counting the number of GFP immunopositive cells throughout the entire retina bilaterally in each fly. The average number of immunopositive cells in the bilateral retina was averaged for each cohort (n=6 per cohort). pH2Av quantification was performed by calculating the percent of pH2Av-positive cells within a 1000x high power field of an anatomically consistent region of mushroom body containing Kenyon cells. The percent of pH2Av-positive cells was averaged for each cohort (n=6 per cohort). H&E and IHC slides were analyzed using a Nikon Eclipse E600 microscope with SPOT software. H&E photos were taken with an Olympus DP25 camera, and IF photos were taken with a Zeiss LSM 800 confocal microscope.

### Statistical analysis

For the climbing assay, chemotherapy cohorts were compared to the H_2_O vehicle control cohort using the Whitney-Mann U test and Bonferroni correction for multiple testing. A two-sided study-wide α level of 0.05 was considered significant. Since there were 9 comparisons (3 different chemotherapeutic agents at 3 concentrations compared to control) per timepoint, a corrected p value threshold of 0.00555 was used to determine significance. For the taste memory assay, the chemotherapy cohorts were compared to the H_2_O vehicle control cohort using a repeated measures 2-way ANOVA with Dunnett’s multiple comparison test. P values <0.05 were considered significant. For the histologic and immunohistochemical studies, chemotherapy cohorts were compared to the H_2_O vehicle control cohort using the Whitney-Mann U test and Bonferroni correction for multiple testing. With a 2-sided study-wide α level of 0.05 for significance and 3 comparisons per timepoint per study (3 concentrations of cisplatin compared to control), a corrected p value threshold of 0.0166 was used to determine significance.

## Acknowledgements

The elav antibodies were obtained from the Developmental Studies Hybridoma Bank created by the National Institute of Child Health and Human Development (National Institutes of Health) and maintained by the Department of Biology at the University of Iowa. The GFP antibody was obtained from the University of California, Davis/National Institutes of Health NeuroMab Facility.

## Data Availability

All study data are included in the article.

## Author Contributions Statement

MT and MBF conceptualized and designed the study. MT performed the experiments. Analysis of the data was performed by MT with statistics consultation by DAM. MT, HB, VN, and CAZ prepared the figures. MT drafted the manuscript. All authors contributed to the critical revision of the manuscript. MBF supervised the study. All authors approved of the manuscript’s submission.

